# Constraint on boric acid resistance and tolerance evolvability in *Candida albicans*

**DOI:** 10.1101/2024.01.04.574193

**Authors:** Yana Syvolos, Ola E. Salama, Aleeza C. Gerstein

**Author notes:** Corresponding author: **Aleeza C. Gerstein**.

## Abstract

Boric acid is a broad-spectrum antimicrobial used to treat vulvovaginal candidiasis when patients relapse on the primary azole drug fluconazole. *Candida albicans* is the most common cause of vulvovaginal candidiasis, colloquially referred to as a "vaginal yeast infection". Little is known about the propensity of *C. albicans* to develop BA resistance or tolerance (the ability of a subpopulation to grow slowly in high levels of drug). We evolved 96 replicates from eight diverse *C. albicans* strains to increasing BA concentrations to test the evolvability of BA resistance and tolerance. Replicate growth was individually assessed daily, with replicates passaged when they had reached an optical density consistent with exponential growth. Many replicates went extinct quickly. Although some replicates could grow in much higher levels of BA than the ancestral strains, evolved populations isolated from the highest terminal BA levels (after 11 weeks of passages) surprisingly showed only modest growth improvements and only at low levels of BA. No large increases in resistance or tolerance were observed in the evolved replicates. Overall, our findings illustrate that there may be evolutionary constraints limiting the emergence of BA resistance and tolerance, which could explain why it remains an effective treatment for recurrent yeast infections.

## Introduction

Vulvovaginal candidiasis (VVC, colloquially "vaginal yeast infection") is an infection of the lower genital tract primarily caused by *Candida albicans* (Sobel 2016). It will affect ∼75% of women at least once during their lifetime (Rosati et al. 2020). Although the majority of people respond well to over-the-counter oral or topical azole treatments, up to 9% will experience recurrent VVC (RVVC), which is defined as at least three symptomatic VVC episodes per year (van Schalkwyk et al. 2015; Sobel 2016; Pappas et al. 2016). Although some risk factors for recurrence have been identified, approximately 50% of the time there is no obvious factor (San Juan Galán et al. 2023). Somewhat surprisingly, acquired drug resistance has been observed in response to maintenance therapy with the first-line drug treatments fluconazole (Marchaim et al. 2012) and clotrimazole (Arastehfar et al. 2021), though does not seem to be the driving factor in the majority of cases. However, many isolates of the second-most common species implicated in RVVC, *Nakaseomyces glabrata* (formerly *Candida glabrata*), have intrinsic resistance to azole drugs (Hassan et al. 2021), and the incidence of non-albicans species may be increasing in prevalence. In addition to drug resistance, fungal drug tolerance, the ability of a subpopulation to grow slowly at inhibitory drug concentrations, has also been potentially implicated in clinical drug failure in other contexts (Delarze and Sanglard 2015; Gerstein et al. 2016; Rosenberg et al. 2018; Berman and Krysan 2020; Levinson et al. 2021). *In vitro* studies have shown that variation in antifungal drug tolerance is independent of variation in resistance (Rosenberg et al. 2018; Salama and Gerstein 2022), and increases in tolerance can also evolve independently of increases in resistance (Gerstein and Berman 2020; Kukurudz et al. 2022a; Todd et al. 2023; Yang et al. 2023).

In some jurisdictions, intravaginal suppositories of boric acid (BA) are used as an alternative treatment for RVVC when first-line treatment fails or when the infection is caused by *N. glabrata* or other intrinsically azole-resistant species (Pappas et al., 2016; van Schalkwyk et al., 2015). Previous studies that measured the susceptibility of *C. albicans* isolates to boric acid have found little variation among isolates consistent with little to no intrinsic resistance (Otero et al. 1999; Romano et al. 2005; De Seta et al. 2009; Salama and Gerstein 2022) and concentrations of 4-10 mg/mL BA inhibit *C. albicans* at rates of 97.2-100% (De Seta et al. 2009; Kalkanci et al. 2012; Larsen et al. 2018). Despite its use as a broad-spectrum and well-tolerated antiseptic against a wide breadth of microbial species and infection types for over a hundred years, BA’s mechanism of action remains unclear. BA is the form of boron present at physiological pH. In *C. albicans*, BA has shown to inhibit virulence factors such as biofilm and hyphae formation (De Seta et al. 2009; Pointer et al. 2015; Salama and Gerstein 2022). In *S. cerevisiae* BA has been implicated in inhibition of oxidative metabolism, and disruption of translational control and amino acid metabolism (Uluisik et al. 2011b; Tsednee et al. 2020). The potential role of boron in cellular growth is unknown. A single study in *S. cerevisiae* found that low concentrations of boron are required, as yeast cells could not divide under acute boron deficiency (Bennett et al. 1999). Although *S. cerevisiae* and *C. albicans* are separated by ∼300 million years of evolution (Hedges et al. 2015), they are commonly grown under similar conditions, indicating common growth requirement; if boron is required for *S. cerevisiae* growth, it may well also be required for *C. albicans*. Directly testing this is surprisingly difficult, as trace amounts of boron are found in all substrates (Schubert 2011).

To test whether *C. albicans* populations evolved to BA would acquire drug resistance or drug tolerance, and possibly gain insight into the mode of action of BA, we conducted the first BA experimental evolution study. The earliest *C. albicans* laboratory evolution study continuously evolved six replicates of an initially susceptible *C. albicans* population to increasing fluconazole concentrations for ∼330 generations (Cowen et al. 2000), finding that all populations increased in resistance. Similar evolution experiments conducted in other species or to other drugs have also since found that resistance readily evolved (e.g., *Candida parapsilosis*: fluconazole, voriconazole, and posaconazole (Papp et al. 2018)*; Candida auris*: fluconazole (Bing et al. 2020)). Evolution experiments conducted in a single drug concentration have recently revealed that the level of drug influences evolutionary trajectory. Two new studies in *C. albicans* found that higher concentrations of fluconazole, i.e, above the minimum inhibitory concentration (MIC_50_), selected for drug tolerance, while resistance arose at lower drug concentrations (Todd et al. 2023; Yang et al. 2023). We hence reasoned that conducting an evolution experimental study in BA should similarly be of utility. We evolved 12 replicates from each of eight different *C. albicans* strain backgrounds to increasing concentrations of BA for up to 26 transfers. We found that even after adjusting the protocol to increase drug concentration very slowly, many replicates went extinct quickly. Furthermore, although some replicates were able to grow in much higher levels of BA than the ancestral strains, evolved populations isolated from the highest terminal BA levels surprisingly only exhibited small growth improvements at low levels of BA, and did not exhibit increased resistance or tolerance. Our results suggest there may be evolutionary constraints limiting the emergence of BA resistance and tolerance, which could explain the rare occurrence of BA resistance clinically (Iavazzo et al. 2011).

## Materials and Methods

### Strains

We selected eight *Candida albicans* strains that capture some of the genotypic breadth of the species: SC5314 (Lockhart et al. 2002); FH1 (Fonzi and Irwin 1993); T101 (Odds et al. 2007); P87, GC75, P78048, P75016, and P76055 (Wu et al. 2007). These strains have been previously characterized genomically (Hirakawa et al. 2015) and were used in experimental evolution studies in fluconazole (Gerstein and Berman 2020; Todd et al. 2023; Yang et al. 2023) and posaconazole (Kukurudz et al. 2022). We refer to these eight strains collectively as the ancestral strains.

### Broth Microdilution Assay

The minimum inhibitory concentration (MIC) of each ancestral strain was measured following the Clinical and Laboratory Standards Institute M27-A2 guidelines (Sheehan et al. 2004) for other fungal drugs with minor modifications. Briefly, strains were treaked from frozen stocks maintained at -70 °C onto Sabouraud dextrose agar (SDA) plates and incubated for 48 h at 37 °C. A loopful of colonies was suspended in 200 μL phosphate-buffered saline (PBS) and standardized to an optical density reading at 600 nm (OD_600_) of 0.01 in 1 mL of PBS. 5 μL of the standardized culture was transferred to a 96-well plate containing 0, 0.2, 0.4, 0.8, and 1 mg/mL of boric acid (BA) in RPMI 1640 (10.4 % w/v RPMI 1640 powder, 1.5 % w/v dextrose, 1.73 % w/v MOPS, adjusted to pH 7 with NaOH tablets; we refer to this media throughout as RPMI). The concentration of BA required to inhibit 50% of growth (MIC_50_) relative to wells containing just RPMI was determined from average OD_600_ readings of 4 technical replicates after 24 h.

### Initial Boric Acid Evolution Experiment

Ancestral strains were streaked from frozen stock onto SDA plates and incubated overnight at 37 °C. A loopful of colonies was transferred to 200 μL PBS, and standardized in PBS to OD_600_ 0.01. Twelve replicates ("ancestral replicates") were then initiated from each ancestral strain by transferring 5 μL of standardized culture into 195 μL RPMI in the twelve wells of the first row of a 96-well plate (i.e., each strain had its own plate). Plates were incubated overnight at 37 °C, and then 5 μL of overnight culture from each well was transferred into a well containing RPMI + 0.4 mg/mL BA in the second row. The plates were incubated statically at 37 °C and taken out every 24 h for an OD_600_ reading. When OD_600_ reached 0.3 (corresponding to visual turbidity in the medium and indicative of logistic growth), 100 μL of culture was transferred to fresh RPMI + 0.8 mg/mL BA, for a final concentration of 0.4 mg/mL BA, and returned to the incubator for continued daily measurement. When the culture again reached OD_600_ 0.3, 100 μL was transferred into fresh RPMI + 1.6 mg/mL BA, for a final concentration of 0.8 mg/mL BA. The cultures were always transferred twice into the same drug concentration, followed by a transfer into medium containing double the previous amount. Wells that did not reach OD_600_ 0.3 after eight days were considered extinct. Drug-evolved replicates were frozen down every second transfer i.e., before increasing the drug concentration. 53% of the replicates (51 of 96) were lost by the fifth transfer (t05) which was into RPMI + 1.6 mg/mL BA, and the remaining 45 replicates went extinct by the seventh transfer, into RPMI + 3.2 mg/mL BA.

### Succeeding Boric Acid Evolution Experiment

After the initial evolution experiment, the protocol was modified in an effort to increase the propensity of replicates that were able to survive growth in higher levels of BA. We refer to this as the succeeding evolution experiment; this linguistic choice is twofold as the experiment was the second iteration and was ultimately effective for achieving our goal of continuing to passage a larger number of replicates to a higher level of boric acid. The succeeding evolution experiment was initiated by inoculating the evolved replicates that were frozen at -70 °C during the initial evolution experiment after two transfers in RPMI + 0.8 mg/mL BA (i.e., t4). The freezer culture was thawed and 2 µL of culture was inoculated into 200 µL RPMI + 0.8 mg/mL BA, and grown overnight at 37 °C. A single replicate (ancestral strain P87, R6) went extinct after the second transfer of the initial evolution experiment (into 0.4 mg/mL, t2, Figure S1b) and was excluded.

To reduce the extinction rate, we reduced the incremental increase of BA concentration. After one overnight in RPMI + 0.8 mg/mL BA, surviving replicates were sequentially grown in RPMI + 1.0 mg/mL, 1.25 mg/mL, 1.5 mg/mL, 1.75 mg/mL, 2 mg/mL, 3 mg/mL, and 4 mg/mL BA. After RPMI + 1.5 mg/mL BA was reached, the number of transfers was changed from two transfers per drug concentration to five transfers. Also after the last transfer in RPMI + 1.5 mg/mL BA, we simultaneously transferred surviving replicates to wells that contained RPMI + 1.75 mg/mL BA and wells that contained RPMI + 2 mg/mL BA. Replicates of five strains (PGC75, P76055, T101, SC5314 and FH1) were unable to grow in RPMI + 2 mg/mL BA within eight days and were classified as group A strains, and the transfer into RPMI + 2 mg/mL BA was discarded. The majority (22 out of 24) of replicates from the other three strains (P87, P78048 and P75016) grew in RPMI + 2 mg/mL BA within eight days and were classified as group B strains, and the transfer into RPMI + 1.75 mg/mL was discarded. As in the initial experiment, cells were frozen down after the last transfer of each drug concentration. No replicates survived growth in RPMI + 4 mg/mL BA. Given the variation among strains in extinction trajectories (i.e., the number of replicates that survived through transfers in different BA concentrations), we chose to focus our phenotypic assessments on a subset of strains. For a balanced design, two strains from each group (Group A: FH1 and GC75; Group B: P75016 and P78048) were chosen. Four ancestral and four evolved replicates isolated from the terminal BA concentration for each strain were examined (FH1: 1.75 mg/mL, GC75: 2 mg/mL, P75016: 3 mg/mL, P78048: 3 mg/mL; Table 1).

**Table 1.** List of replicates used in phenotypic analysis. Evolved replicates were isolated from their terminal boric acid (BA) concentration. Group A: FH1 and GC75; Group B: P75016 and P78048.

| Strain | Transfer | Ancestral Replicates | Evolved Replicates | [BA] (mg/ml) |
| --- | --- | --- | --- | --- |
| FH1 | t18 | R4, R5, R8 | R1, R2, R3, R4 | 1.75 |
| GC75 | t23 | R2, R3, R7, R8 | R2, R3, R4, R8 | 2 |
| P75016 | t23 | R7, R8, R9, R10 | R3, R4, R7, R8 | 3 |
| P78048 | t23 | R3, R5, R6, R7 | R6, R7, R8, R9 | 3 |

### Disk Diffusion Assay

Ancestral and evolved replicates from the four chosen strains were grown from frozen by streaking to single colony on SDA plates and incubated for 48 h at 37 °C. A loopful of colonies from each replicate was then suspended in 200 μL of PBS and standardized to OD_600_ 0.01 in 1 mL of PBS. 100 μL of standardized culture was spread in duplicates onto 15 mL Muller Hinton (MH) plates using sterile glass beads. Plates were left to dry before a 5 mg boric acid disk was placed in the center of each plate. BA disks were prepared by transferring 10 μL of preheated 500 mg/mL BA in dimethyl sulfoxide (DMSO) stock to blank antimicrobial disks (6 mm, Fisher Scientific). MH plates were incubated at 37 °C for 48 h. Photographs of each plate on a light box were taken at 48 h. As previously described, ImageJ was used to edit photographs (Salama and Gerstein 2022) before quantification of drug susceptibility and tolerance by the R package *diskImageR* (Gerstein et al. 2016). Briefly, photographs were cropped to a uniform size, the colours were inverted, and brightness and contrast were adjusted with consistent parameters across all images to maximize the contrast between the white disk and the black background. Susceptibility was measured as RAD_20_, the radius where 20% reduction of growth occurs, while tolerance was measured as FoG_20_, the fraction of growth above RAD_20_ ^(^Gerstein et al. 2016^)^.

### Growth in the evolutionary drug concentrations

Ancestral and evolved replicates were grown from frozen by streaking onto SDA plates and then incubated for 48 h at 37 °C. A colony from each replicate was resuspended in 200 μL of PBS and standardized to OD_600_ of 0.01 in 1 mL of PBS. 200 μL of RPMI or RPMI + BA stock were added to each well of a 96-well round bottom plate, for final concentrations of 0 mg/mL, 0.2 mg/mL, 0.4 mg/mL, 0.8 mg/mL, 1 mg/mL, 1.25 mg/mL, 1.5 mg/mL, and 1.75 mg/mL of BA, and 5 μL of standardized culture was added to each well. Plates were covered with a Breathe-Easy sealing membrane (Electron Microscopy Sciences, PA, United States) and incubated shaking at 37 °C for 72 h. OD_600_ measurements were taken at the end of the incubation period. The average OD_600_ between three technical replicates from each ancestral and evolved replicate was used for analysis and visualization.

### Growth Rate Assay

Two growth rate assays, in RPMI and RPMI + 1.25 mg/mL BA, were conducted. To initiate each experiment, ancestral and evolved replicates were grown from frozen by streaking onto SDA plates and incubated for 48 h at 37 °C. A loop full of colonies from each replicate was then resuspended in 200 μL of PBS and standardized to OD_600_ of 0.01 in 1 mL of PBS. 5 µL of standardized culture was added into each well of a 96-well round bottom plate containing 150 µL of RPMI or RPMI + 1.25 mg/mL BA. The plates were covered with a Breathe-Easy sealing membrane (Electron Microscopy Sciences, PA, United States) and incubated in a Synergy H1 microplate spectrophotometer (Agilent Technologies, CA, United States) at 37°C for 72 h shaking continuously. OD_600_ measurements were taken every 15 minutes. From each well, the maximal growth rate was calculated as the spline with the highest slope using a custom R script written by Dr. Richard Fitzjohn. The average growth rate between three technical replicates was used for analysis and visualization.

### Statistical analyses

Details on statistical analyses are included throughout the text where relevant. The data and code required to reproduce all data analyses and visualizations is available here: https://github.com/acgerstein/boric_acid_evolution/

## Results

### Trajectory of *C. albicans* evolution in BA

We evolved twelve replicate lines from eight ancestral genotypes (SC5314, FH1, T101, P87, GC75, P78048, P75016, and P76055; total of 96 replicates) to increasing concentrations of boric acid (BA). The ancestral BA minimum inhibitory concentration (MIC_50_) for T101 was 0.4 mg/mL, and 0.8 mg/mL for all other strains. An initial evolution experiment was conducted where 53% of replicates went extinct by the fifth transfer, into 1.6 mg/mL BA, and all replicates were extinct by the seventh transfer into 3 mg/mL (see Methods for additional details). This prompted us to initiate a second ("succeeding" experiment). In the succeeding experiment, the incremental increase in BA concentration was decreased and the number of transfers at each drug concentration was increased to five starting at 1.5 mg/mL BA. These experimental changes increased the number of replicates that survived to higher drug concentrations. At transfer 14, the last transfer in 1.5 mg/mL BA, the remaining evolved replicates were split into two groups: replicates from group A strains (GC75, P76055, T101, SC5314 and FH1) were transferred into 1.75 mg/mL BA; replicates from group B strains (P87, P78048 and P75016) were transferred into 2 mg/mL BA (Figure 1). Group A strains went extinct in 2 and 3 mg/mL BA, while group B replicates went extinct at 3 and 4 mg/mL BA.

**Figure 1.**
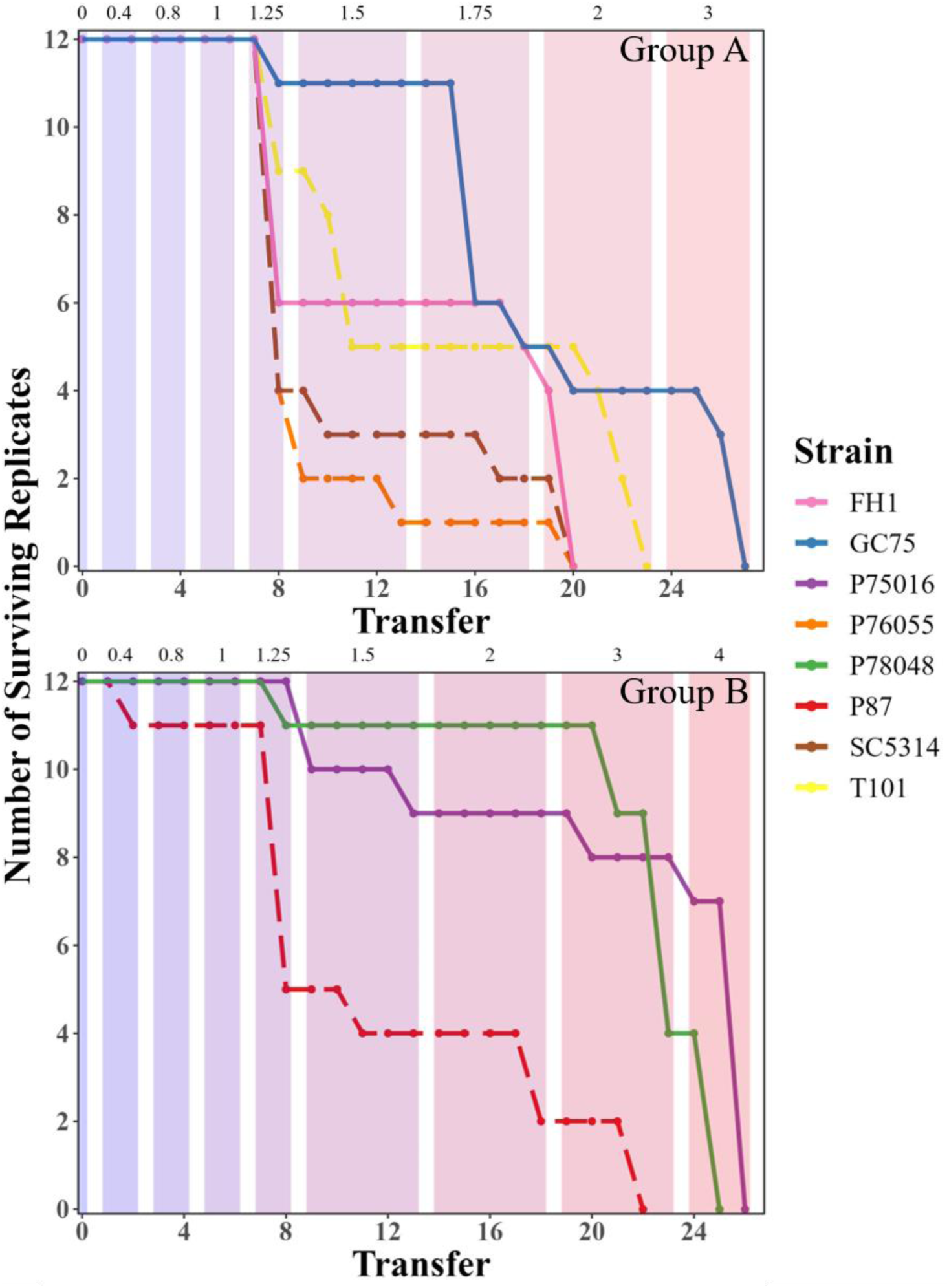
Boric acid evolution experiment. Eight *C. albicans* strains grown from frozen on SDA at 37 °C were transferred through increasing concentrations of BA in RPMI medium. The numbers at the top of each panel indicate the BA concentration. Note that after RPMI + 1.25 mg/mL BA, the number of transfers in each drug concentration increased from two to five. Replicates were frozen down at the last transfer of each drug concentration. Solid lines represent the strains selected for phenotypic testing.

The ancestral genotype significantly influenced the terminal drug concentration that was reached by evolved replicates. Group A replicates first began going extinct after the seventh transfer (1.25 mg/mL BA) when 43% of all Group A replicates died (Figure 1). The remaining Group A replicates went extinct in 2 mg/mL BA, with the exception of four replicates of GC75 that survived to 3 mg/mL BA before extinction (Figure 1). By contrast, only 20% of Group B replicates were extinct after the seventh transfer (1.25 mg/mL BA). The remaining P87 replicates went extinct at or prior to 3 mg/mL while seven P76016 and four P78048 replicates survived to be transferred into 4 mg/mL BA prior to extinction (Figure 1). Hence the terminal drug concentration reached strongly depended on ancestral strain background.

Somewhat surprisingly, the number of days between transfers was variable and was not obviously linked to genotype or drug concentration. Replicates were always transferred when the OD_600_ reached a minimum of 0.3. In the succeeding evolution experiment, initially there was a linear increase in the number of days between transfers. It took only one day for all replicates to grow and be transferred from 0.8 mg/mL BA (ancestral MIC_50_) into 1 mg/mL BA, and two days from 1 mg/mL BA into 1.25 mg/mL BA (Figure S3). All of the replicates also took the same amount of time, three days, to be transferred in 1 mg/mL BA. However, at higher BA concentrations, the time between transfers into the same or different concentration varies greatly, from 1 to 8 days, between replicates (Figure S3).

### Phenotypic analysis of BA evolved replicates

We focused the phenotypic analysis on the terminal replicates, from four strains that were isolated from BA concentrations of 1.75 mg/mL BA (FH1, group A), 2 mg/mL BA (GC75, group A) and 3 mg/mL BA (P75016 and P78048, group B). This corresponds to BA concentrations between 2.19× to 3.75× above their initial MIC_50_ (Table 1). Surprisingly, when measured with the disk diffusion assay, there was no significant change in resistance between ancestral and evolved replicates of P75016, P78048, or FH1 (Figure 2A, statistical results from Welch two sample t-tests in Table 2) while the evolved replicates of GC75 had a significant decrease in resistance compared to their ancestors, albeit with a small effect size (Figure 2A, Table 2). There was also no significant difference in tolerance between the ancestral and evolved replicates from any of the strains (Figure 2B).

**Figure 2.**
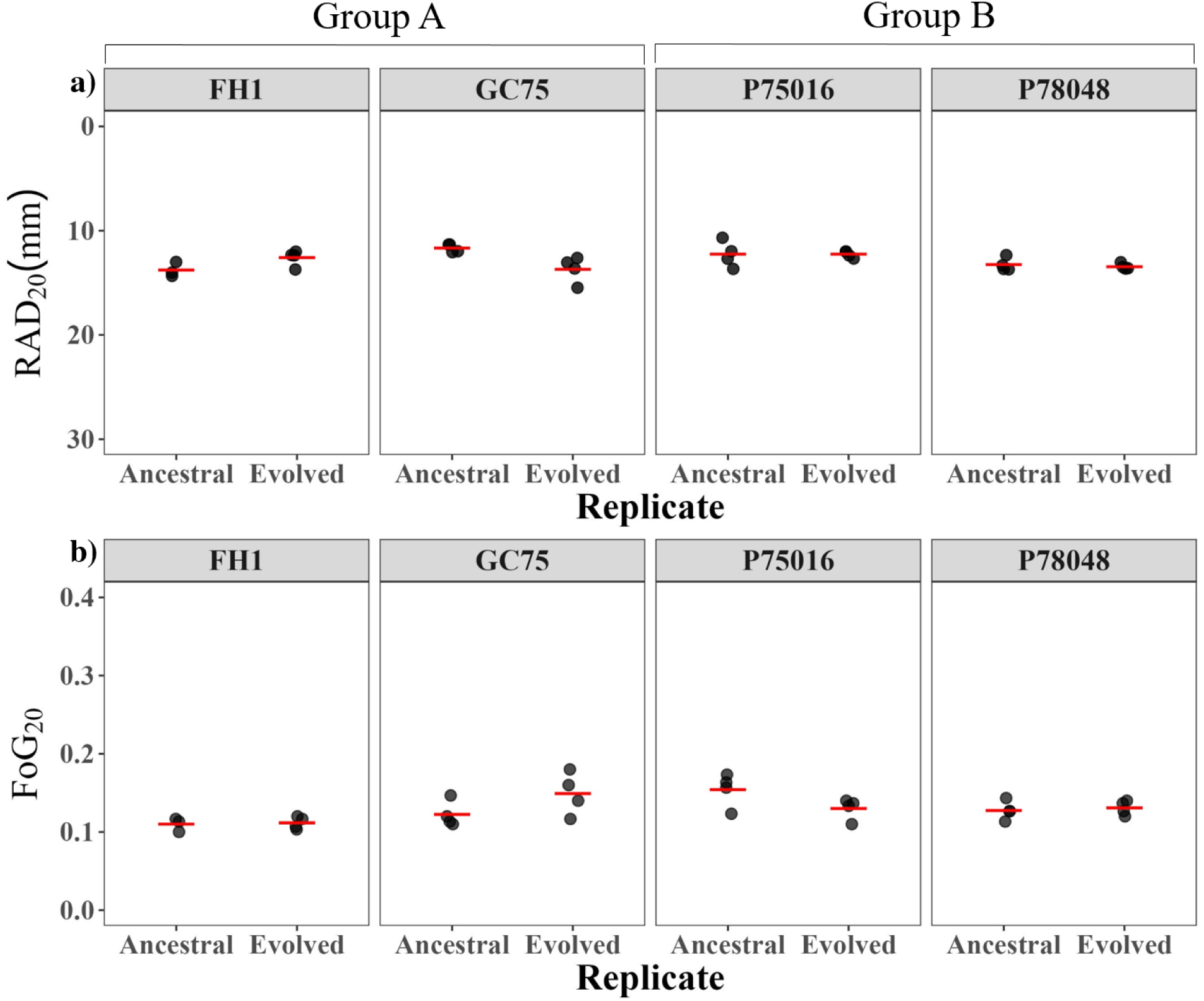
Variation in susceptibility (a) and tolerance (b) between ancestral and evolved replicates. The y-axis for susceptibility is reversed so that more resistant replicates (i.e., replicates that grew closer to the drug disk) are towards the top. Red lines indicate the mean.

**Table 2.**
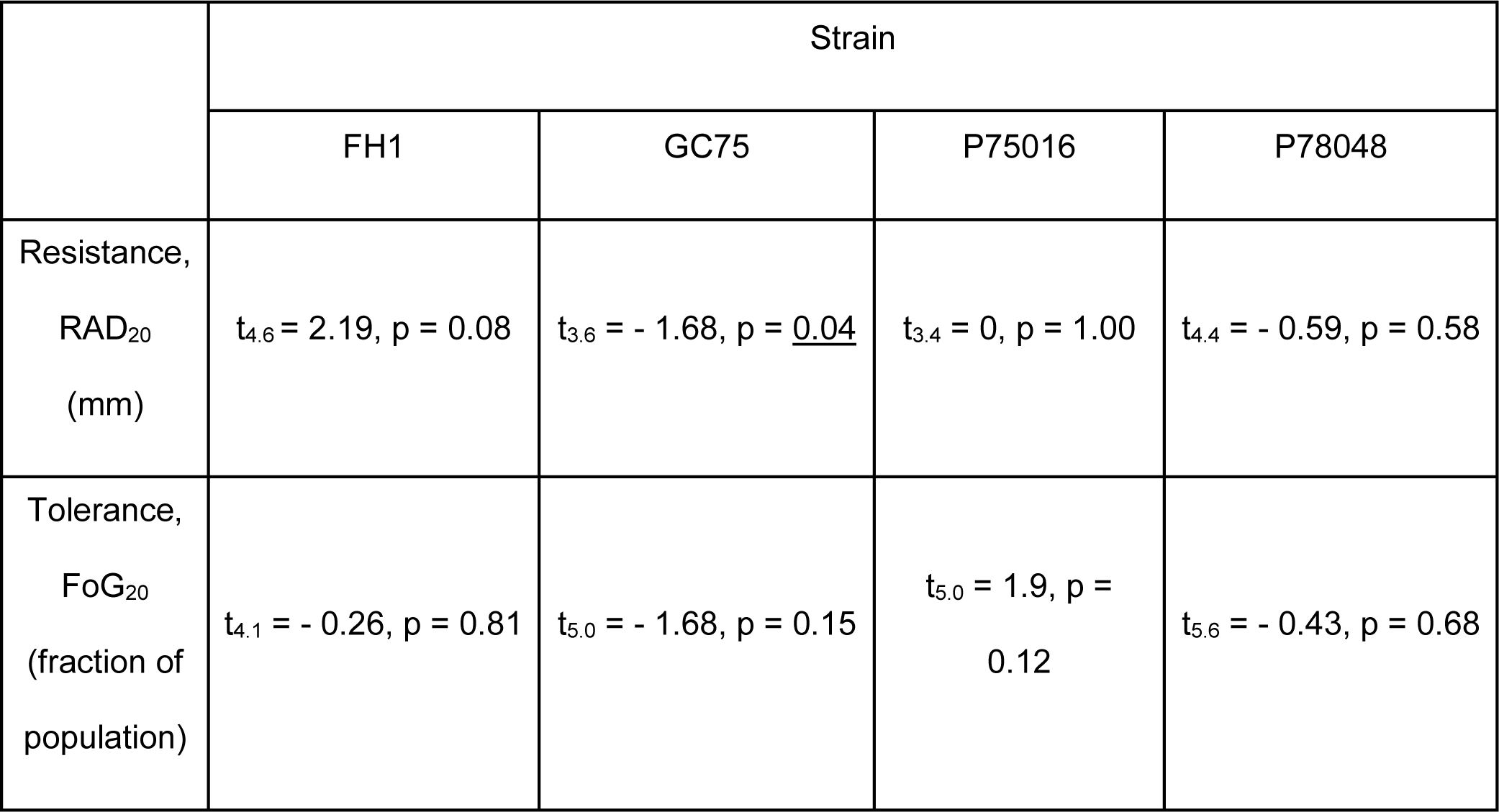
Welch two-sample t-tests comparing resistance and tolerance between ancestral and evolved replicates. Significant p-values are underlined.

In addition to drug resistance and tolerance, we also examined whether there were growth improvements (assessed as optical density at 600 nm) in seven different BA concentrations after 24 and 72 h of growth. Evolved replicates from all strains grew better at levels of BA up to the ancestral MIC_50_ (i.e., at 0.2 mg/mL, 0.4 mg/mL, and 0.8 mg/mL, Figure 3, statistical results from Welch two sample t-tests in Table 3). However, the growth advantages were largely restricted to low levels of BA that were ∼1.5-2× lower than the terminal BA concentration (Figure 3, Table 3). Note that the timing of an advantage (24 h vs 72 h) and the drug levels where improvements were observed were somewhat idiosyncratic, e.g., GC75 had an improvement in 72 h but not 24 h in 1 mg/mL BA and 1.25 mg/mL but not 0.8 mg/mL. In addition, the majority of evolved replicates from three strains (FH1, GC75, P75016) grew better than the ancestral strains at 24 and 72 h in RPMI with no drug; growth of evolved replicates from P78048 was similar to the ancestral strain. Overall, despite the lack of resistance or tolerance phenotypes, all strains exhibited growth improvements albeit at BA levels much lower than those from which they were isolated, and in the absence of BA.

**Figure 3.**
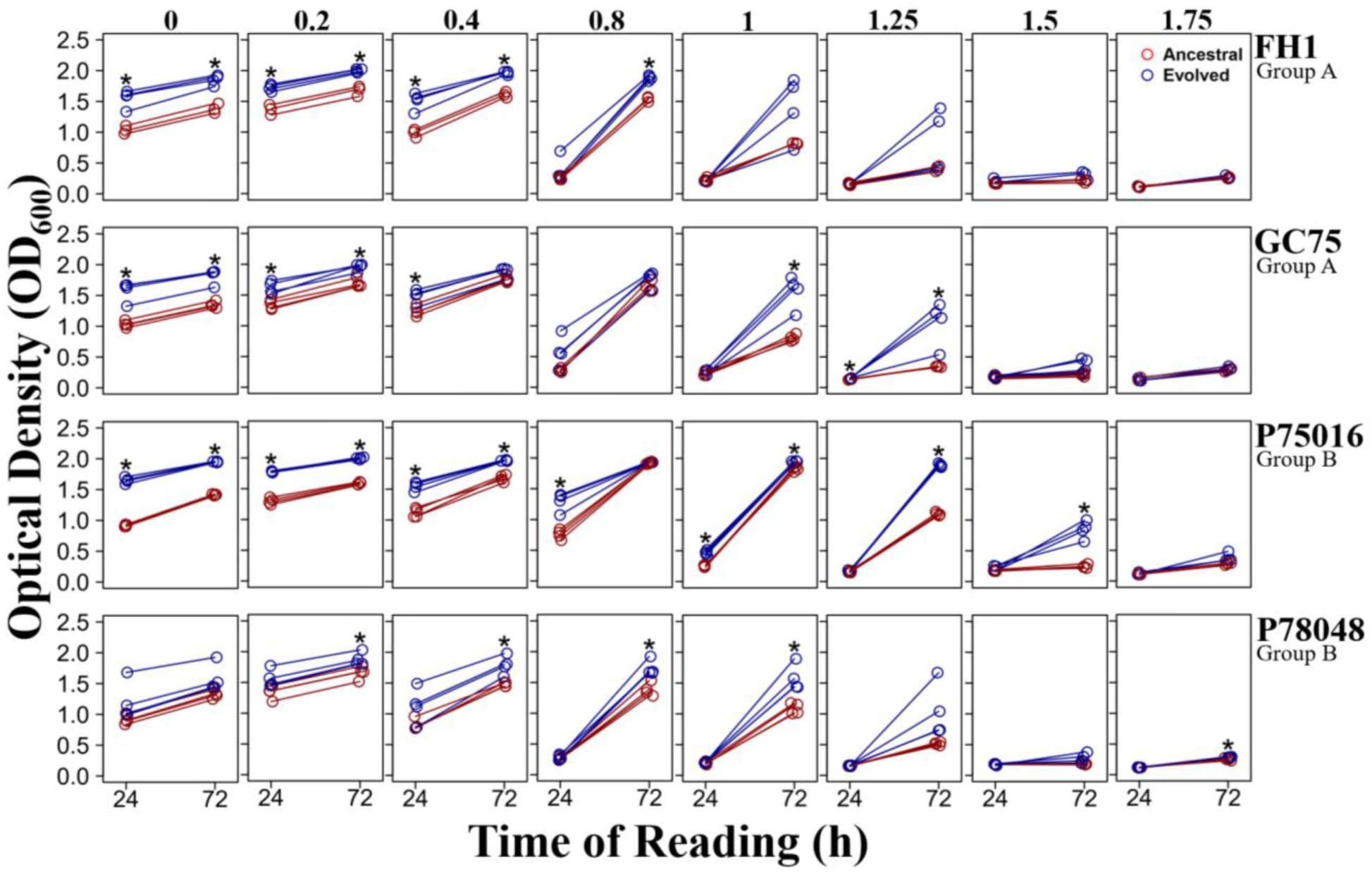
Fitness of ancestral and evolved replicates grown in RPMI and increasing concentrations of BA. Numbers at the top represent BA concentration. Optical density was measured at 24 h and 72 h. Ancestral replicates are shown in red, evolved replicates are shown in dark blue. Stars indicate statistical significance in a t-test comparing ancestral and evolved replicates at one time point (24 h or 72 h of growth).

**Table 3.** Welch two-sample t-tests to compare ancestral and evolved fitness in different BA concentrations. Significant p-values are underlined.

| [BA]<br>(mg/mL) | Time<br>(h) | Strain |  |  |  |
| --- | --- | --- | --- | --- | --- |
|  |  | FH1 | GC75 | P75016 | P78048 |
| 0 | 24 | $t_{4.3} = -6.22, p = \underline{0.003}$ | $t_{3.6} = -6.25, p = \underline{0.005}$ | $t_{3.7} = -27.78, p < \underline{0.001}$ | $t_{3.3} = -1.75, p = 0.170$ |
| | 72 | $t_{4.5} = -7.77, p = \underline{0.001}$ | $t_{4.0} = -7.01, p = \underline{0.002}$ | $t_{5.6} = -70.26, p = \underline{0.001}$ | $t_{3.5} = -2.17, p = 0.106$ |
| 0.2 | 24 | $t_{3.7} = -3.44, p = \underline{0.029}$ | $t_{5.4} = -4.41, p = \underline{0.005}$ | $t_{3.3} = -17.83, p = \underline{0.002}$ | $t_{5.5} = -2.48, p = 0.051$ |
| | 72 | $t_{3.3} = -4.03, p = \underline{0.023}$ | $t_{5.8} = -5.80, p = \underline{0.001}$ | $t_{5.8} = -30.87, p < \underline{0.001}$ | $t_{6.0} = -3.06, p = \underline{0.022}$ |
| 0.4 | 24 | $t_{4.6} = -2.67, p = 0.050$ | $t_{5.3} = -3.16, p = \underline{0.023}$ | $t_{6.0} = -8.30, p = \underline{0.001}$ | $t_{3.6} = -2.03, p = 0.121$ |
| | 72 | $t_{3.3} = -4.40, p = \underline{0.018}$ | $t_{5.1} = -2.41, p = 0.060$ | $t_{3.2} = -10.05, p = \underline{0.002}$ | $t_{3.3} = -3.96, p = \underline{0.024}$ |
| 0.8 | 24 | $t_{5.2} = -0.41, p = 0.700$ | $t_{3.1} = -2.22, p = 0.112$ | $t_{4.5} = -6.30, p = \underline{0.002}$ | $t_{3.4} = -0.66, p = 0.552$ |
| | 72 | $t_{4.5} = -10.87, p = \underline{0.001}$ | $t_{5.4} = -1.85, p = 0.119$ | $t_{4.3} = -1.51, p = 0.202$ | $t_{5.8} = -4.11, p = \underline{0.007}$ |
| 1 | 24 | $t_{3.5} = 0.98, p = 0.391$ | $t_{5.6} = -0.39, p = 0.715$ | $t_{4.0} = -9.84, p = \underline{0.001}$ | $t_{5.0} = -2.51, p = 0.054$ |
| | 72 | $t_{5.6} = -1.21, p = 0.275$ | $t_{3.2} = -5.53, p = \underline{0.010}$ | $t_{4.6} = -5.33, p = \underline{0.004}$ | $t_{4.0} = -4.28, p = \underline{0.013}$ |
| 1.25 | 24 | $t_{3.9} = 0.176, p = 0.869$ | $t_{6.0} = -3.68, p = \underline{0.010}$ | $t_{5.9} = -0.55, p = 0.603$ | $t_{5.6} = -0.032, p = 0.976$ |
| | 72 | $t_{3.2} = -1.47, p = 0.232$ | $t_{3.0} = -3.97, p = \underline{0.029}$ | $t_{6.0} = -39.81, p < \underline{0.001}$ | $t_{3.0} = -2.40, p = 0.095$ |
| 1.5 | 24 | $t_{3.7} = -2.29, p = 0.089$ | $t_{4.1} = -0.07, p = 0.946$ | $t_{3.4} = -1.53, p = 0.214$ | $t_{3.5} = 0.39, p = 0.719$ |
| | 72 | $t_{3.6} = -2.03, p = 0.120$ | $t_{3.4} = -2.35, p = 0.091$ | $t_{3.3} = -8.01, p = \underline{0.003}$ | $t_{3.1} = -2.58, p = 0.081$ |
| 1.75 | 24 | $t_{5.2} = 0.339, p = 0.748$ | $t_{5.8} = 0.285, p = 0.786$ | $t_{3.2} = -0.386, p = 0.724$ | $t_{4.3} = -1.21, p = 0.288$ |
| | 72 | $t_{5.3} = -0.97, p = 0.375$ | $t_{5.7} = -1.47, p = 0.196$ | $t_{3.3} = -2.70, p = 0.066$ | $t_{3.5} = -3.56, p = \underline{0.030}$ |

To gain additional precision on potential differences between ancestral and evolved replicates, we measured growth rate in the absence of BA and in 1.25 mg/mL BA (a level of drug slightly above the ancestral MIC_50_ where the majority of evolved replicates had increased growth at 72h hours). In the absence of BA, evolved replicates on average had a higher growth rate than the ancestral replicates in FH1 (t_3.8_ = -7.95, p = 0.002), GC75 (t_3.0_= -4.92, p = 0.016), P75016, (t_4.7_ = -17.71, p < 0.001) but not P78048 (t_3.0_ = -1.82, p = 0.17) (Figure 4A). Evolved replicates also exhibited an increased growth rate on average in 1.25 mg/mL BA in GC75 (t_3.8_ = -3.66, p = 0.023), P78048 (t_3.0_ = -4.09, p = 0.026) and P75016 (t_3.5_ = -5.19, p = 0.011), but not from FH1 (t_3.0_ = -2.21, p = 0.11) (Figure 4B).

**Figure 4.**
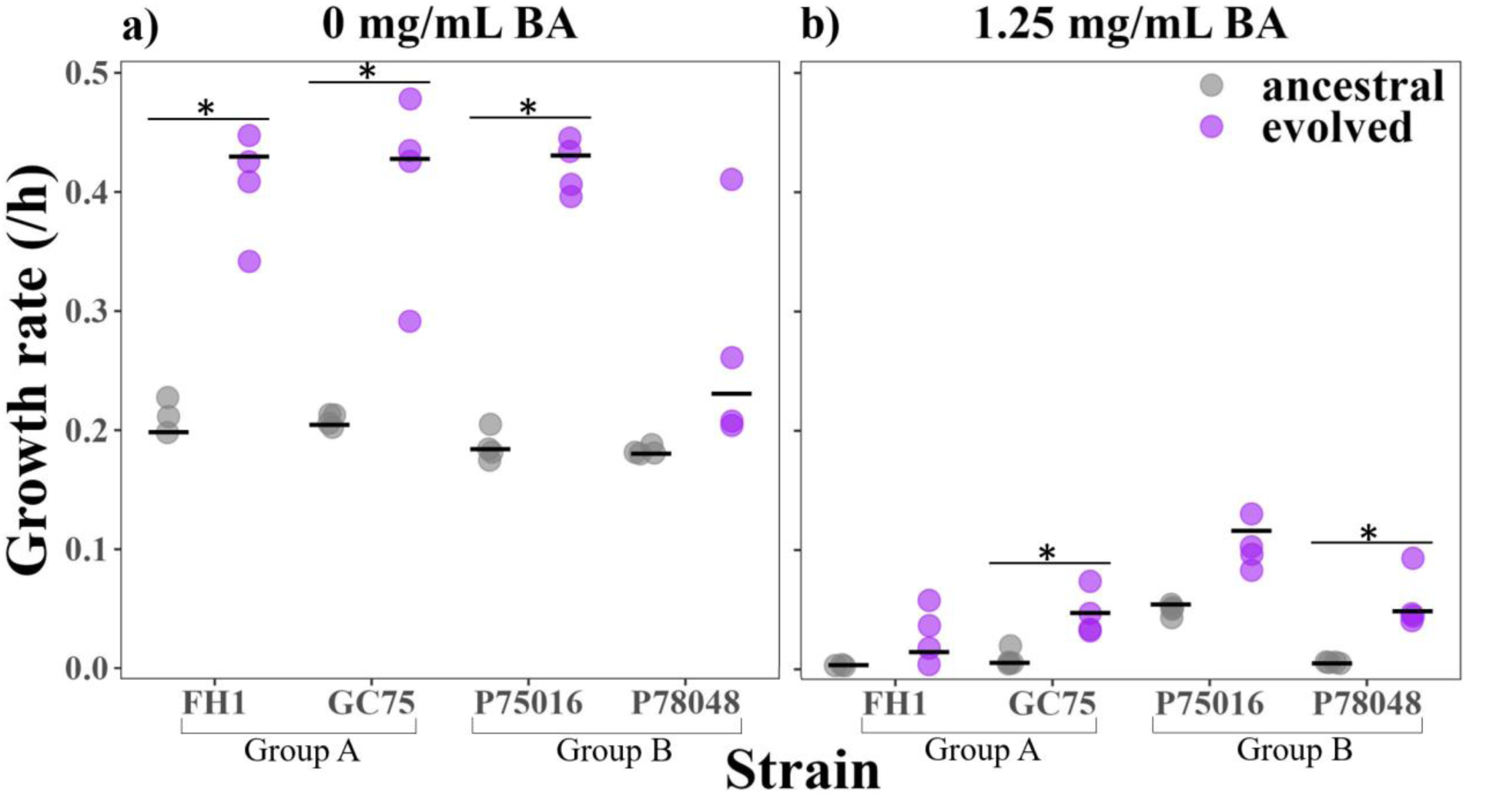
Growth rate was measured from 4 ancestral and 4 evolved replicates from FH1, GC75, P7016, and P78048 *C. albicans* strains in a) RPMI and b) RPMI + 1.25 mg/mL BA. Optical density was recorded every 15 minutes in a plate reader with constant shaking and incubation at 37 °C. Each point represents the mean of three technical replicates. Growth rate was calculated as the spline with the highest slope using a custom R script. Stars indicate statistical significance in a t-test comparing growth rates of ancestral and evolved replicates.

The overarching pattern through analysis of replicates evolved to BA through *in vitro* evolution was many replicates going extinct at low BA concentrations (1 - 1.5 mg/mL BA). Replicates that survived to the highest levels of drug (1.75 - 4 mg/mL BA) experienced evolutionary changes that resulted in improved growth, but only at no/low levels of drug and to a very small increase in drug beyond the ancestral MIC_50_ level.

## Discussion

We evolved 96 replicates from eight genotypically diverse *C. albicans* strains to increasing boric acid (BA) concentrations to assess the evolvability of BA resistance and tolerance. We found that many replicates quickly went extinct in drug levels just above the ancestral minimum inhibitory concentration (MIC_50_). Furthermore, although some surviving replicates were passaged in drug concentrations up to 3.75× ancestral MIC_50_, growth increases were nearly all restricted to growth in the absence of BA, low BA concentrations up to the ancestral MIC_50_, and within a very narrow range above the ancestral MIC_50_. Surprisingly, the highest BA concentration at which evolved replicates exhibited improved growth was 2.2× lower than the BA concentration that they were isolated from.

The fact that the observed small improvement in growth was restricted to drug concentrations much lower than the drug concentration replicates were isolated from is distinct from what has been observed in the majority of published *in vitro* evolution studies of antifungal drug resistance. Previous experiments that used a similar experimental design, where the level of drug was continually increased (“stepwise experimental evolution”), typically report that resistance often increased to drug levels that far exceed the ancestral MIC_50_ in both azoles and other drug classes: 256× in *C. albicans* strains evolved to fluconazole, Cowen et al. 2000; 8× to 256× in fluconazole evolved *C. tropicalis*, Paul et al. 2020; 8× in *S. cerevisiae* evolved to fluconazole, Anderson et al. 2003; 128× fluconazole, 258× voriconazole, and 1032× posaconazole evolved *C. parapsilosis*, Papp et al. 2020; *C. parapsilosis* evolved to echinocandins (4× caspofungin, 8× anidulafungin, and 8× micafungin), Papp et al. 2018; 512× in *S. cerevisiae* evolved to amphotericin B, Anderson et al. 2009. Here, despite isolation from BA concentrations above the ancestral MIC_50_, none of the terminal evolved replicates increased in resistance compared to their ancestor; in fact, GC75 showed a decrease in resistance, albeit small, suggesting that there must be significant barriers to the acquisition of BA resistance in *C. albicans*. Despite the perceived success of previous experimental evolution studies, there are two important caveats to note. In many published experiments, the end-point was deliberately chosen to be when an increased MIC_50_ was achieved or included a final experimental step that only selected resistant isolates for further analysis. Few studies have provided a description of isolates that went extinct or those that did not increase in resistance (Gerstein and Sethi 2022). The second is a pervasive publication bias against null results. It may be that other *in vitro* experimental studies also did not result in resistance increases, yet that they have remained unpublished.

In the majority of previous studies, the resistant phenotype was maintained for several generations after passage in drug free media, indicating it was genetically encoded (Cowen et al. 2000; Papp et al. 2018, 2020; Ksiezopolska et al. 2021; Bergin et al. 2022). However, Paul et al. (2020) observed that five of their fluconazole resistant *C. tropicalis* isolates, with MICs ≤ 32 mg/L, reverted to low MIC (1.0 mg/L) after only 5 subcultures in fluconazole free media, and the remaining four *C. tropicalis* isolates, with MICs ≥ 128 mg/L reverted after 20 subcultures. This rapid phenotypic conversion is typically ascribed to chromosomal aneuploidies, which can be rapidly lost in the absence of selection. We do not think that is the likely explanation here, as only limited growth in the absence of drug (i.e., 48 h of growth in RPMI) was typically conducted prior to drug exposure for phenotypic analysis. The transient ability of the replicate populations to grow in high levels of BA may be epigenetically regulated. At higher BA concentrations (beyond 1.25 mg/mL BA), the time between transfers into the same or different BA concentration varied greatly, from 1 to 8 days. Recent studies have shown that epigenetic changes in chromatin remodelling and histone acetylation or deacetylation can contribute to resistance in *C. albicans* to azole and echinocandin antifungal drugs <u>(Wurtele et al. 2010; Li et al. 2015; Liu and Myers 2017)</u>. As the mode of action for BA remains elusive, it is difficult to speculate how the terminal evolved populations were all able to reach an OD_600_ of 0.3 in much higher levels of drug than they had an observed growth benefit in. Boric acid seems to damage *C. albicans’* cytoplasmic membrane over time (De Seta et al. 2009). Uluisik et al. (2011a) also suggested that in *S. cerevisiae* boron exerts its toxic effect through activation of the general amino acid control system and inhibition of protein synthesis. Future work will examine gene expression and epigenetic regulation including histone modifications and DNA methylation patterns to gain insight into how *C. albicans* populations that have not acquired bona fide resistance or tolerance can grow in high BA levels when the drug concentration is slowly increased.

Boric acid resistance evolution may be restricted by fitness trade-offs. Similar to boric acid, clinical resistance to amphotericin B remains extremely rare despite 50 years of monotherapeutic use (Gallis et al. 1990; Vincent et al. 2013). Vincent et al. (2013) found that mutations that confer resistance to amphotericin B in *C. albicans* create a variety of severe and costly cellular stresses that decrease the cell’s ability to tolerate external stresses from the host. A potentially relevant study examined *S. cerevisiae* replicates that were acquired following growth in 12 mM copper, yet all had an MIC**_50_** below that level (Gerstein et al. 2015). Copper is a known essential micronutrient for *S. cerevisiae* that is toxic at high concentrations. A single study in *S. cerevisae* found that they were unable to divide in the absence of BA (Bennett et al. 1999). If this is the case for *C. albicans*, then they would be unable to evolve to block BA uptake entirely, critically limiting the pathways to adaptation. Boron is said to be present in trace amounts everywhere in nature (Schubert 2011), thus it may be that its essential function in *C. albicans* has gone unnoticed because boron deficiency is never observed naturally.

The difficulty of *C. albicans* populations to evolve resistance and/or tolerance *in vitro* is consistent with the rare occurrence of clinical resistance (Iavazzo et al. 2011). This general constraint of BA resistance evolution contrasts with the repeated observation of azole resistance arising during clinical use and potentially has significant clinical implications (White et al. 2002). Future studies need to examine the evolvability of BA resistance and tolerance in other VVC-causing species, such as *N. glabrata*, as BA is often used to treat infections caused by this species. Overall, our findings show that continuous *in vitro* exposure of *C. albicans* replicates to increasing concentrations of BA *in vitro* selected for a small window of improved growth in low levels of BA, but did not select for clinically meaningful BA-resistance or BA-tolerance.

## Acknowledgements

A.C.G acknowledges the support of the CIFAR Azrieli Global Scholars Program. This work was additionally supported by a Discovery Grant to A.C.G. from the Natural Sciences and Engineering Research Council of Canada (NSERC). O.E.S was supported by a University of Manitoba Graduate Fellowship (UMGF). We thank Dr. Lee Flippin and Dr. Nediljko Budisa for thoughtful conversations about boric acid and Dr. Lee Flippin, in particular, for his encouragement of our work in this space.

## Supplementary Material

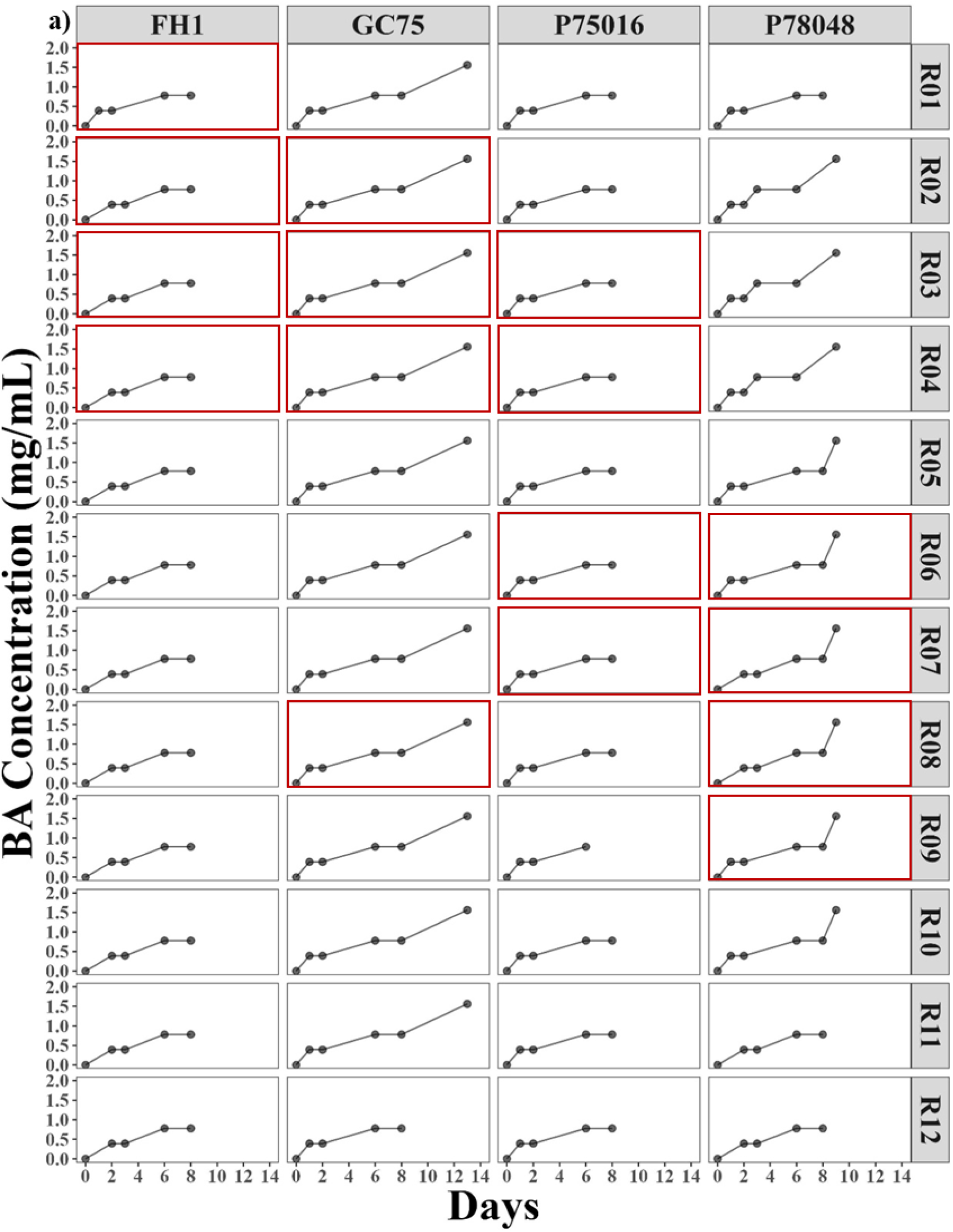

## References

Anderson, J.B., Sirjusingh, C., Parsons, A.B., Boone, C., Wickens, C., Cowen, L.E., and Kohn, L.M. 2003. Mode of selection and experimental evolution of antifungal drug resistance in Saccharomyces cerevisiae. Genetics 163(4): 1287–1298.

Anderson, J.B., Sirjusingh, C., Syed, N., and Lafayette, S. 2009. Gene expression and evolution of antifungal drug resistance. Antimicrob. Agents Chemother. 53(5): 1931–1936.

Arastehfar, A., Carvalho, A., Houbraken, J., Lombardi, L., Garcia-Rubio, R., Jenks, J.D., Rivero-Menendez, O., Aljohani, R., Jacobsen, I.D., Berman, J., Osherov, N., Hedayati, M.T., Ilkit, M., Armstrong-James, D., Gabaldón, T., Meletiadis, J., Kostrzewa, M., Pan, W., Lass-Flörl, C., Perlin, D.S., and Hoenigl, M. 2021. Aspergillus fumigatus and aspergillosis: From basics to clinics. Stud. Mycol. 100: 100115.

Bennett, A., Rowe, R.I., Soch, N., and Eckhert, C.D. 1999. Boron stimulates yeast (Saccharomyces cerevisiae) growth. J. Nutr. 129(12): 2236–2238.

Bergin, S.A., Zhao, F., Ryan, A.P., Müller, C.A., Nieduszynski, C.A., Zhai, B., Rolling, T., Hohl, T.M., Morio, F., Scully, J., Wolfe, K.H., and Butler, G. 2022. Systematic Analysis of Copy Number Variations in the Pathogenic Yeast Candida parapsilosis Identifies a Gene Amplification in RTA3 That is Associated with Drug Resistance. MBio 13(5): e0177722.

Berman, J., and Krysan, D.J. 2020. Drug resistance and tolerance in fungi. Nat. Rev. Microbiol. 18(6): 319–331.

Bing, J., Hu, T., Zheng, Q., Muñoz, J.F., Cuomo, C.A., and Huang, G. 2020. Experimental Evolution Identifies Adaptive Aneuploidy as a Mechanism of Fluconazole Resistance in Candida auris. Antimicrob. Agents Chemother. 65(1). doi:10.1128/AAC.01466-20.

Cowen, L.E., Sanglard, D., Calabrese, D., Sirjusingh, C., Anderson, J.B., and Kohn, L.M. 2000. Evolution of drug resistance in experimental populations of Candida albicans. J. Bacteriol. 182(6): 1515–1522.

Delarze, E., and Sanglard, D. 2015. Defining the frontiers between antifungal resistance, tolerance and the concept of persistence. Drug Resist. Updat. 23: 12–19.

De Seta, F., Schmidt, M., Vu, B., Essmann, M., and Larsen, B. 2009. Antifungal mechanisms supporting boric acid therapy of Candida vaginitis. J. Antimicrob. Chemother. 63(2): 325– 336.

Fonzi, W.A., and Irwin, M.Y. 1993. Isogenic strain construction and gene mapping in Candida albicans. Genetics 134(3): 717–728.

Gallis, H.A., Drew, R.H., and Pickard, W.W. 1990. Amphotericin B: 30 years of clinical experience. Rev. Infect. Dis. 12(2): 308–329.

Gerstein, A.C., and Berman, J. 2020. Candida albicans Genetic Background Influences Mean and Heterogeneity of Drug Responses and Genome Stability during Evolution in Fluconazole. mSphere 5(3). doi:10.1128/mSphere.00480-20.

Gerstein, A.C., Ono, J., Lo, D.S., Campbell, M.L., Kuzmin, A., and Otto, S.P. 2015. Too much of a good thing: the unique and repeated paths toward copper adaptation. Genetics 199(2): 555–571.

Gerstein, A.C., Rosenberg, A., Hecht, I., and Berman, J. 2016. diskImageR: quantification of resistance and tolerance to antimicrobial drugs using disk diffusion assays. Microbiology 162(7): 1059–1068.

Gerstein, A.C., and Sethi, P. 2022. Experimental evolution of drug resistance in human fungal pathogens. Curr. Opin. Genet. Dev. 76: 101965.

Hassan, Y., Chew, S.Y., and Than, L.T.L. 2021. Candida glabrata: Pathogenicity and Resistance Mechanisms for Adaptation and Survival. J Fungi (Basel) 7(8). doi:10.3390/jof7080667.

Hatwig, C., Balbueno, E.A., Bergamo, V.Z., Pippi, B., Fuentefria, A.M., and Silveira, G.P. 2019. Multidrug-resistant *Candida glabrata* strains obtained by induction of anidulafungin resistance in planktonic and biofilm cells. Braz. J. Pharm. Sci. 55: e18025. Universidade de São Paulo, Faculdade de Ciências Farmacêuticas. [accessed 11 October 2023].

Hedges, S.B., Marin, J., Suleski, M., Paymer, M., and Kumar, S. 2015. Tree of life reveals clock-like speciation and diversification. Mol. Biol. Evol. 32(4): 835–845.

Hirakawa, M.P., Martinez, D.A., Sakthikumar, S., Anderson, M.Z., Berlin, A., Gujja, S., Zeng, Q., Zisson, E., Wang, J.M., Greenberg, J.M., Berman, J., Bennett, R.J., and Cuomo, C.A. 2015. Genetic and phenotypic intra-species variation in Candida albicans. Genome Res. 25(3): 413–425.

Iavazzo, C., Gkegkes, I.D., Zarkada, I.M., and Falagas, M.E. 2011. Boric acid for recurrent vulvovaginal candidiasis: the clinical evidence. J. Womens. Health 20(8): 1245–1255.

Kalkanci, A., Güzel, A.B., Khalil, I.I.J., Aydin, M., Ilkit, M., and Kuştimur, S. 2012. Yeast vaginitis during pregnancy: susceptibility testing of 13 antifungal drugs and boric acid and the detection of four virulence factors. Med. Mycol. 50(6): 585–593.

Ksiezopolska, E., Schikora-Tamarit, M.À., Beyer, R., Nunez-Rodriguez, J.C., Schüller, C., and Gabaldón, T. 2021. Narrow mutational signatures drive acquisition of multidrug resistance in the fungal pathogen Candida glabrata. Curr. Biol. 31(23): 5314–5326.e10.

Kukurudz, R.J., Chapel, M., Wonitowy, Q., Adamu Bukari, A.-R., Sidney, B., Sierhuis, R., and Gerstein, A.C. 2022. Acquisition of cross-azole tolerance and aneuploidy in Candida albicans strains evolved to posaconazole. G3 12(9). doi:10.1093/g3journal/jkac156.

Larsen, B., Petrovic, M., and De Seta, F. 2018. Boric Acid and Commercial Organoboron Products as Inhibitors of Drug-Resistant Candida albicans. Mycopathologia 183(2): 349– 357.

Levinson, T., Dahan, A., Novikov, A., Paran, Y., Berman, J., and Ben-Ami, R. 2021. Impact of tolerance to fluconazole on treatment response in Candida albicans bloodstream infection. Mycoses 64(1): 78–85.

Liu, Z., and Myers, L.C. 2017. Candida albicans Swi/Snf and Mediator Complexes Differentially Regulate Mrr1-Induced MDR1 Expression and Fluconazole Resistance. Antimicrob. Agents Chemother. 61(11). doi:10.1128/AAC.01344-17.

Li, X., Cai, Q., Mei, H., Zhou, X., Shen, Y., Li, D., and Liu, W. 2015. The Rpd3/Hda1 family of histone deacetylases regulates azole resistance in Candida albicans. J. Antimicrob. Chemother. 70(7): 1993–2003.

Lockhart, S.R., Pujol, C., Daniels, K.J., Miller, M.G., Johnson, A.D., Pfaller, M.A., and Soll, D.R. 2002. In Candida albicans, white-opaque switchers are homozygous for mating type. Genetics 162(2): 737–745.

Marchaim, D., Lemanek, L., Bheemreddy, S., Kaye, K.S., and Sobel, J.D. 2012. Fluconazole-resistant Candida albicans vulvovaginitis. Obstet. Gynecol. 120(6): 1407–1414.

Odds, F.C., Bougnoux, M.-E., Shaw, D.J., Bain, J.M., Davidson, A.D., Diogo, D., Jacobsen, M.D., Lecomte, M., Li, S.-Y., Tavanti, A., Maiden, M.C.J., Gow, N.A.R., and d’Enfert, C. 2007. Molecular phylogenetics of Candida albicans. Eukaryot. Cell 6(6): 1041–1052.

Otero, L., Fleites, A., Méndez, F.J., Palacio, V., and Vázquez, F. 1999. Susceptibility of Candida species isolated from female prostitutes with vulvovaginitis to antifungal agents and boric acid. Eur. J. Clin. Microbiol. Infect. Dis. 18(1): 59–61.

Pappas, P.G., Kauffman, C.A., Andes, D.R., Clancy, C.J., Marr, K.A., Ostrosky-Zeichner, L., Reboli, A.C., Schuster, M.G., Vazquez, J.A., Walsh, T.J., Zaoutis, T.E., and Sobel, J.D. 2016. Clinical Practice Guideline for the Management of Candidiasis: 2016 Update by the Infectious Diseases Society of America. Clin. Infect. Dis. 62(4): e1–50.

Papp, C., Bohner, F., Kocsis, K., Varga, M., Szekeres, A., Bodai, L., Willis, J.R., Gabaldón, T., Tóth, R., Nosanchuk, J.D., Vágvölgyi, C., and Gácser, A. 2020. Triazole Evolution of Candida parapsilosis Results in Cross-Resistance to Other Antifungal Drugs, Influences Stress Responses, and Alters Virulence in an Antifungal Drug-Dependent Manner. mSphere 5(5). doi:10.1128/mSphere.00821-20.

Papp, C., Kocsis, K., Tóth, R., Bodai, L., Willis, J.R., Ksiezopolska, E., Lozoya-Pérez, N.E., Vágvölgyi, C., Mora Montes, H., Gabaldón, T., Nosanchuk, J.D., and Gácser, A. 2018. Echinocandin-Induced Microevolution of Candida parapsilosis Influences Virulence and Abiotic Stress Tolerance. mSphere 3(6). Am Soc Microbiol. doi:10.1128/mSphere.00547-18.

Paul, S., Singh, S., Sharma, D., Chakrabarti, A., Rudramurthy, S.M., and Ghosh, A.K. 2020. Dynamics of in vitro development of azole resistance in Candida tropicalis. J Glob Antimicrob Resist 22: 553–561.

Pointer, B.R., Boyer, M.P., and Schmidt, M. 2015. Boric acid destabilizes the hyphal cytoskeleton and inhibits invasive growth of Candida albicans. Yeast 32(4): 389–398.

Romano, L., Battaglia, F., Masucci, L., Sanguinetti, M., Posteraro, B., Plotti, G., Zanetti, S., and Fadda, G. 2005. In vitro activity of bergamot natural essence and furocoumarin-free and distilled extracts, and their associations with boric acid, against clinical yeast isolates. J. Antimicrob. Chemother. 55(1): 110–114.

Rosati, D., Bruno, M., Jaeger, M., Ten Oever, J., and Netea, M.G. 2020. Recurrent Vulvovaginal Candidiasis: An Immunological Perspective. Microorganisms 8(2). doi:10.3390/microorganisms8020144.

Rosenberg, A., Ene, I.V., Bibi, M., Zakin, S., Segal, E.S., Ziv, N., Dahan, A.M., Colombo, A.L., Bennett, R.J., and Berman, J. 2018. Antifungal tolerance is a subpopulation effect distinct from resistance and is associated with persistent candidemia. Nat. Commun. 9(1): 2470.

Salama, O.E., and Gerstein, A.C. 2022. Differential Response of Candida Species Morphologies and Isolates to Fluconazole and Boric Acid. Antimicrob. Agents Chemother. 66(5): e0240621.

San Juan Galán, J., Poliquin, V., and Gerstein, A.C. 2023. Insights and advances in recurrent vulvovaginal candidiasis. PLoS Pathog. 19(11): e1011684.

van Schalkwyk, J., Yudin, M.H., and INFECTIOUS DISEASE COMMITTEE. 2015. Vulvovaginitis: screening for and management of trichomoniasis, vulvovaginal candidiasis, and bacterial vaginosis. J. Obstet. Gynaecol. Can. 37(3): 266–274.

Schubert, D. 2011, April 15. Boron Oxides, Boric Acid, and Borates. John Wiley & Sons, Inc., Hoboken, NJ, USA. doi:10.1002/0471238961.0215181519130920.a01.pub3.

Sheehan DJ, Brown SD, Pfaller MA, Warnock DW, Rex JH, Chaturvedi V, Espinel-Ingroff A, Ghannoum MA, Moore LS, Odds FC, Rinaldi MG, Walsh TJ. 2004. Method for antifungal disk diffusion susceptibility testing of yeasts: approved guideline M27-A2. Clinical and Laboratory Standards Institute, Wayne, PA.

Sobel, J.D. 2016. Recurrent vulvovaginal candidiasis. Am. J. Obstet. Gynecol. 214(1): 15–21.

Todd, R.T., Soisangwan, N., Peters, S., Kemp, B., Crooks, T., Gerstein, A., and Selmecki, A. 2023. Antifungal Drug Concentration Impacts the Spectrum of Adaptive Mutations in Candida albicans. Mol. Biol. Evol. 40(1). doi:10.1093/molbev/msad009.

Tsednee, M., Tanaka, M., Kasai, K., and Fujiwara, T. 2020. Boron-dependent regulation of translation through AUGUAA sequence in yeast. Yeast 37(12): 638–646.

Uluisik, I., Kaya, A., Fomenko, D.E., Karakaya, H.C., Carlson, B.A., Gladyshev, V.N., and Koc, A. 2011a. Boron stress activates the general amino acid control mechanism and inhibits protein synthesis. PLoS One 6(11): e27772.

Uluisik, I., Kaya, A., Unlu, E.S., Avsar, K., Karakaya, H.C., Yalcin, T., and Koc, A. 2011b. Genome-wide identification of genes that play a role in boron stress response in yeast. Genomics 97(2): 106–111.

Vincent, B.M., Lancaster, A.K., Scherz-Shouval, R., Whitesell, L., and Lindquist, S. 2013. Fitness trade-offs restrict the evolution of resistance to amphotericin B. PLoS Biol. 11(10): e1001692.

White, T.C., Holleman, S., Dy, F., Mirels, L.F., and Stevens, D.A. 2002. Resistance mechanisms in clinical isolates of Candida albicans. Antimicrob. Agents Chemother. 46(6): 1704–1713.

Wurtele, H., Tsao, S., Lépine, G., Mullick, A., Tremblay, J., Drogaris, P., Lee, E.-H., Thibault, P., Verreault, A., and Raymond, M. 2010. Modulation of histone H3 lysine 56 acetylation as an antifungal therapeutic strategy. Nat. Med. 16(7): 774–780.

Wu, W., Lockhart, S.R., Pujol, C., Srikantha, T., and Soll, D.R. 2007. Heterozygosity of genes on the sex chromosome regulates Candida albicans virulence. Mol. Microbiol. 64(6): 1587– 1604.

Yang, F., Scopel, E.F.C., Li, H., Sun, L.-L., Kawar, N., Cao, Y.-B., Jiang, Y.-Y., and Berman, J. 2023. Antifungal Tolerance and Resistance Emerge at Distinct Drug Concentrations and Rely upon Different Aneuploid Chromosomes. MBio 14(2): e0022723.

