## Supplementary figures and images for "Constraint on boric acid resistance and tolerance evolvability in *Candida albicans*"

### Supplemental Figure 1a

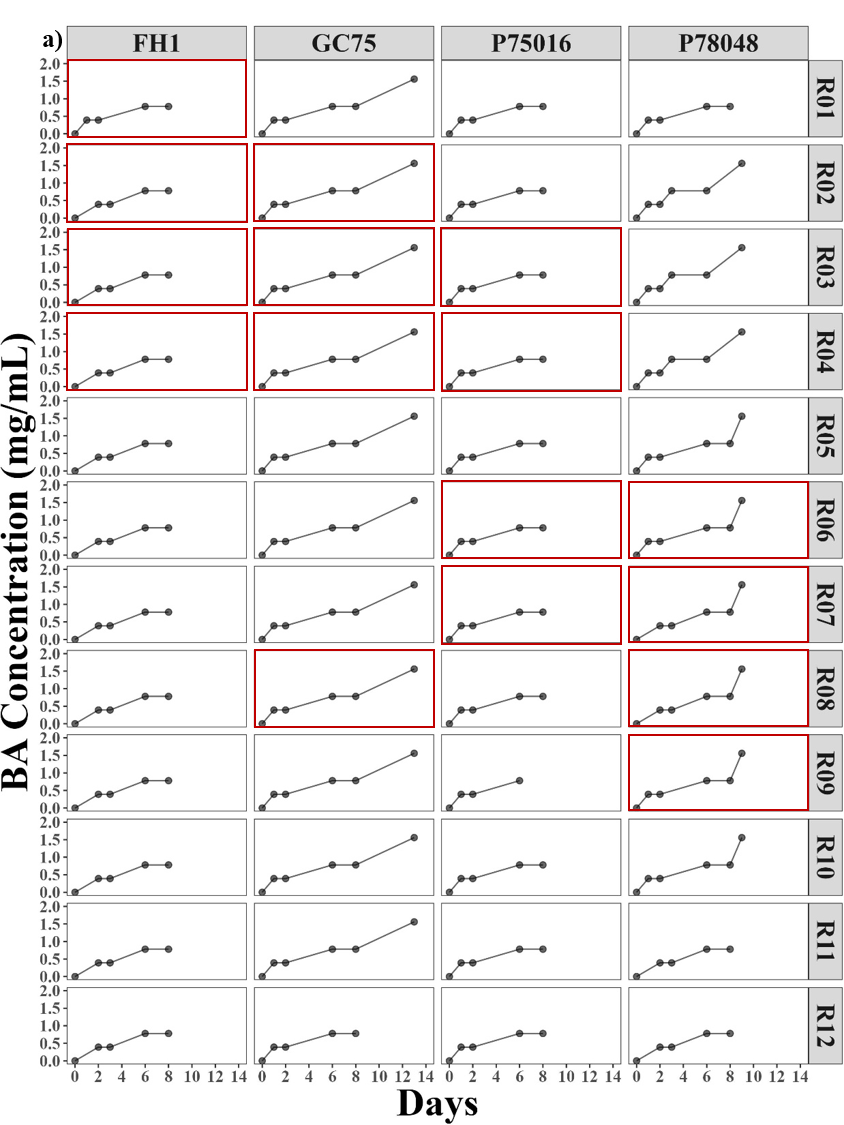

### Supplemental Figure 1a

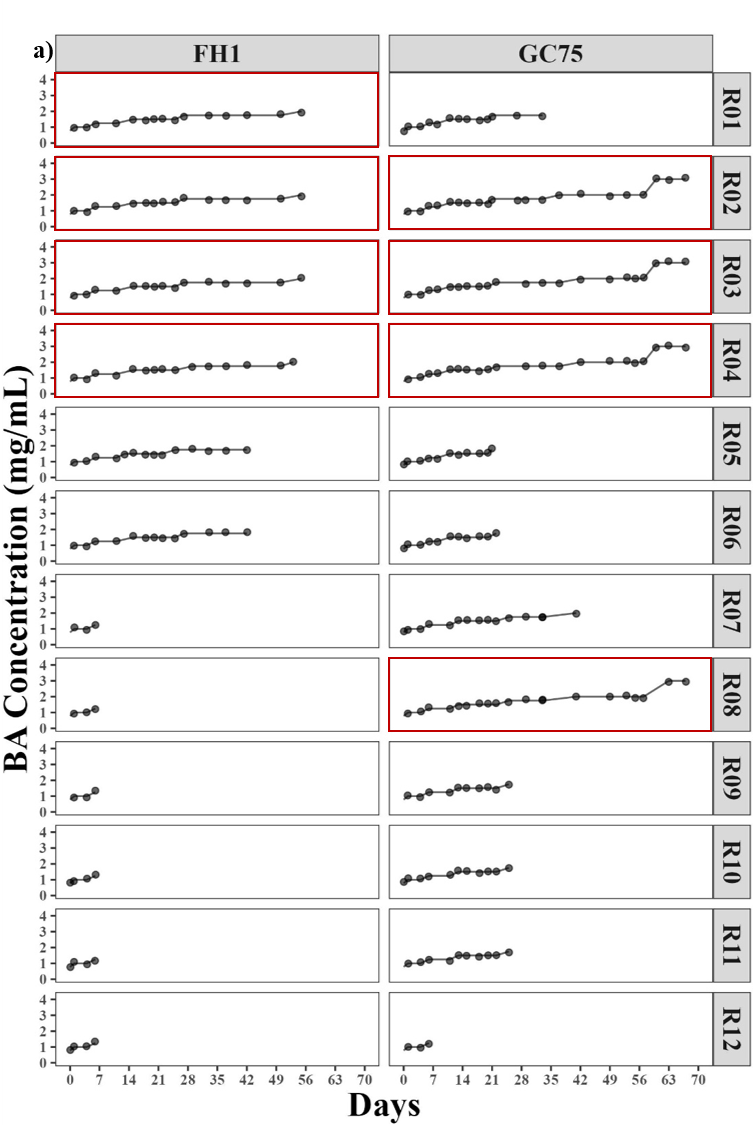

### Supplemental Figure 1b

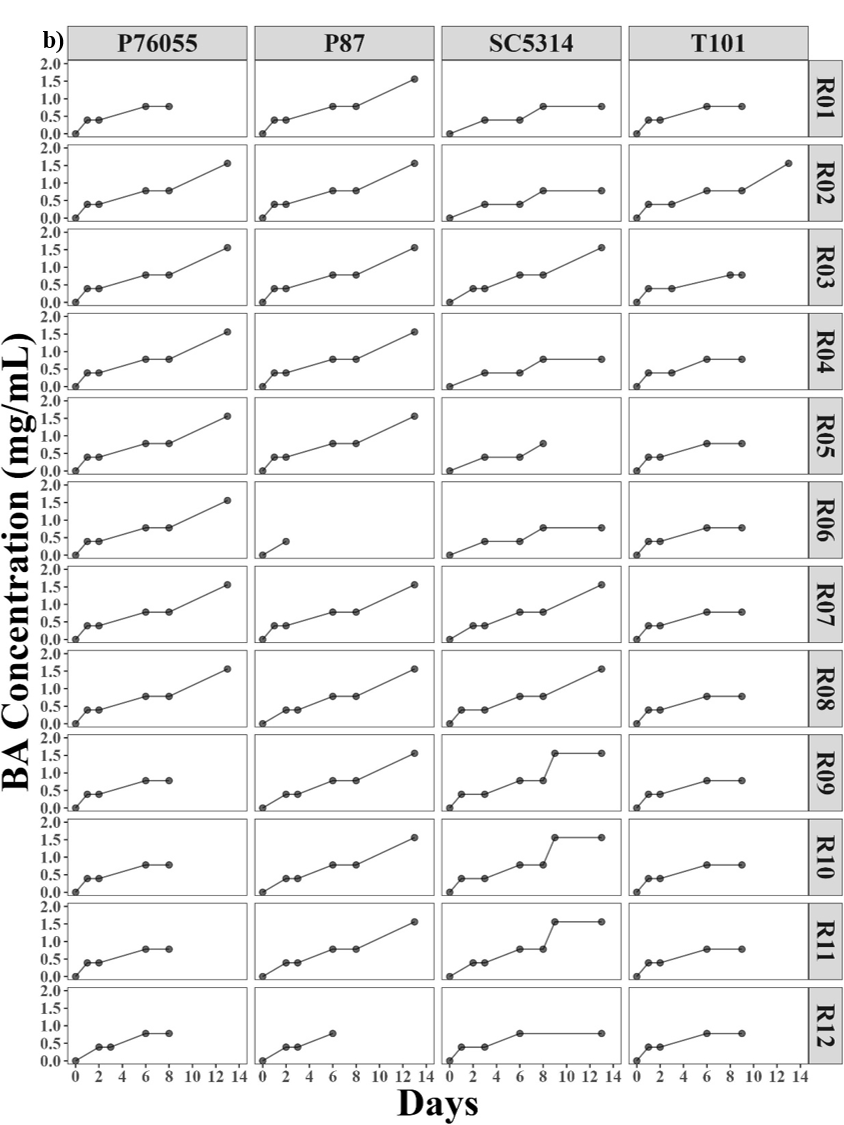

### Supplemental Figure 2b

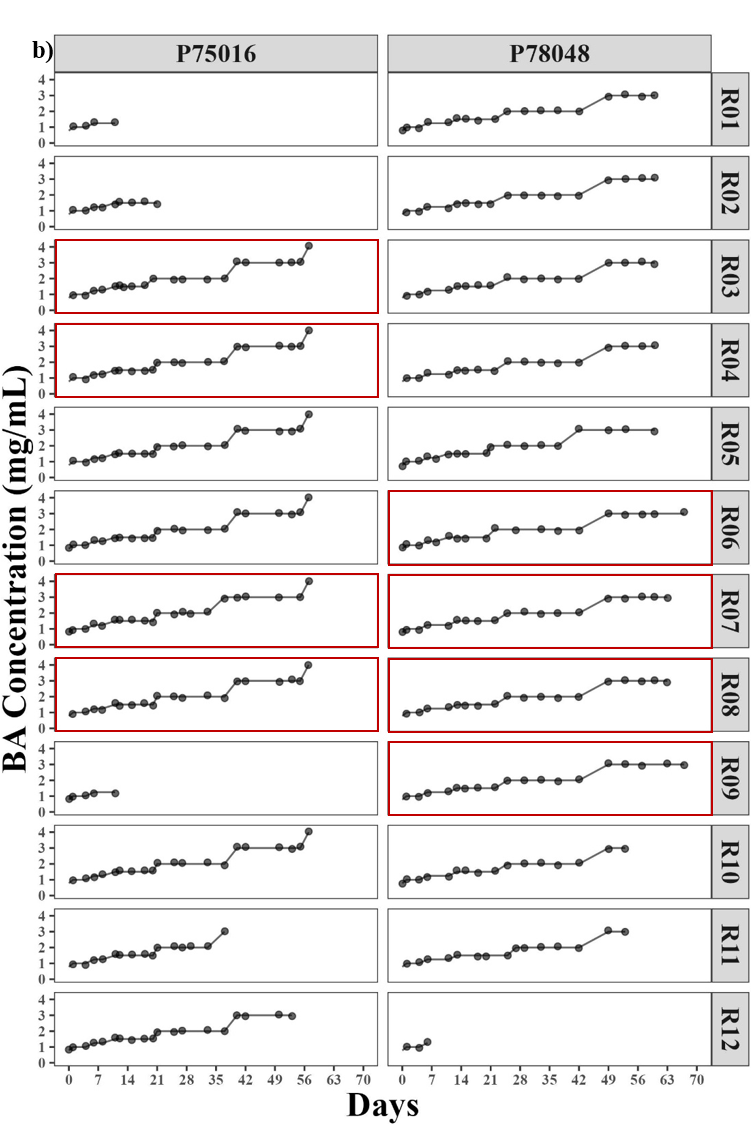

### Supplemental Figure 2c

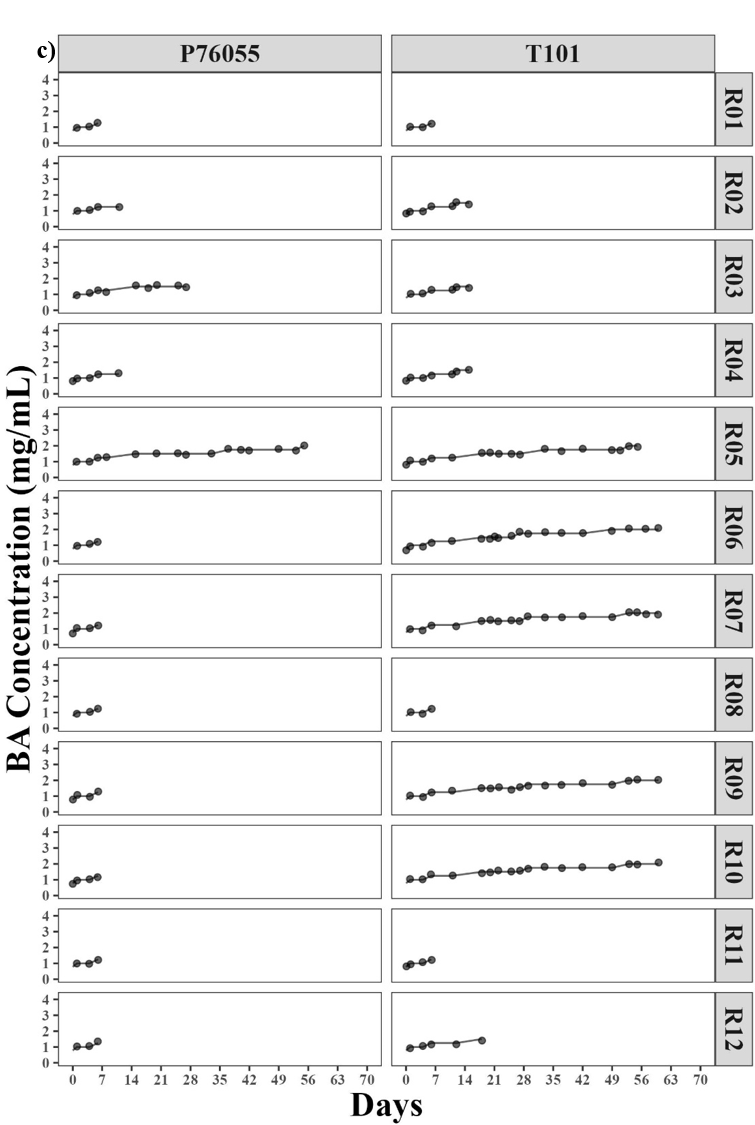

### Supplemental Figure 2d

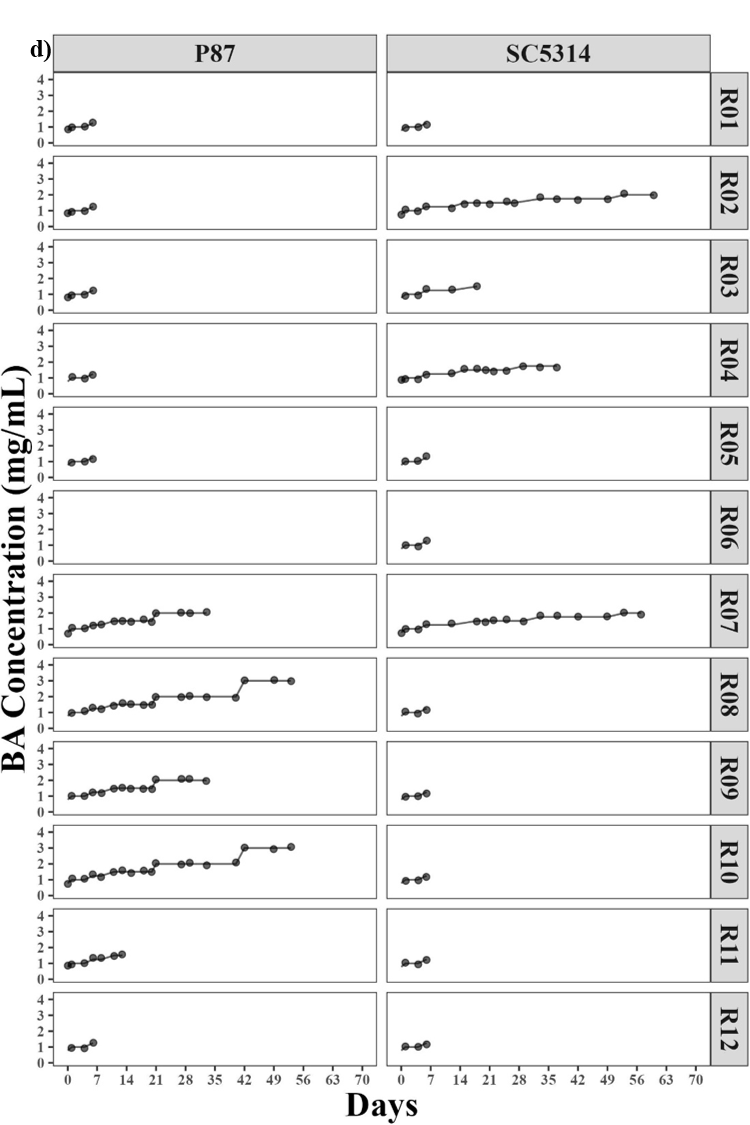
